# Estrogen receptors in human bladder cells regulate innate cytokine responses to differentially modulate uropathogenic *E. coli* colonization

**DOI:** 10.1101/2020.05.27.120089

**Authors:** Ayantika Sen, Anil Kaul, Rashmi Kaul

## Abstract

The bladder epithelial cells elicit robust innate immune responses against urinary tract infections (UTIs) for preventing the bacterial colonization. Physiological fluctuations in circulating estrogen levels in women increase the susceptibility to UTI pathogenesis, often resulting in adverse health outcomes. Dr adhesin bearing *Escherichia coli* (Dr *E. coli*) cause recurrent UTIs in menopausal women and acute pyelonephritis in pregnant women. Dr *E. coli* bind to epithelial cells via host innate immune receptor CD55, under hormonal influence. The role of estrogens or estrogen receptors (ERs) in regulating the innate immune responses in the bladder are poorly understood. In the current study, we investigated the role of ERα, ERβ and GPR30 in modulating the innate immune responses against Dr *E. coli* induced UTI using human bladder epithelial carcinoma 5637 cells (HBEC). Both ERα and ERβ agonist treatment in bladder cells induced a protection against Dr *E. coli* invasion via upregulation of TNFα and downregulation of CD55 and IL10, and these effects were reversed by action of ERα and ERβ antagoinsts. In contrast, the agonist-mediated activation of GPR30 led to an increased bacterial colonization due to suppression of innate immune factors in the bladder cells, and these effects were reversed by the antagonist-mediated suppression of GPR30. Further, siRNA-mediated ERα knockdown in the bladder cells reversed the protection against bacterial invasion observed in the ERα positive bladder cells, by modulating the gene expression of TNFα, CD55 and IL10, thus confirming the protective role of ERα. We demonstrate for the first time a protective role of nuclear ERs, ERα and ERβ but not of membrane ER, GPR30 against Dr *E. coli* invasion in HBEC 5637 cells. These findings have many clinical implications and suggest that ERs may serve as potential drug targets towards developing novel therapeutics for regulating local innate immunity and treating UTIs.

**Highlights:**

- Estrogen receptor (ER) subtypes regulate the gene expression of innate immune molecules, CD55, TNFα and IL10 in human bladder epithelial cells impacting the bacterial colonization by Dr *E. coli*.
- Activation of nuclear ER subtypes, ERα and ERβ, upregulate the gene expression of proinflammatory cytokine, TNFα, but downregulate the gene expression of anti-inflammatory cytokine, IL10, and Dr *E. coli* colonization receptor, CD55, thus leading to efficient bacterial clearance in human bladder cells.
- In contrast, activation of the membrane ER subtype, GPR30, shows opposite effects to ERα and ERβ that were mediated on TNFα, IL10 and CD55 gene expression, thus leading to impaired bacterial clearance of Dr *E. coli* in human bladder cells.
- ER subtypes can serve as potential drug candidates for designing new therapies to boost or modulate the local immunity in the human bladder preventing the establishment of *E. coli* infections.

## 1. Introduction

Urinary tract infection (UTI) may involve the infection of any part of the urinary tract. Ascending UTIs start with the infection of the urethra causing urethritis, may progress into the bladder to cause cystitis. In worst cases, if the host immunity is compromised in the bladder, the infection can advance to the kidneys causing acute or chronic pyelonephritis. Dr fimbriae bearing *Escherichia coli (*Dr *E. coli)*-induced recurrent UTI in menopausal women or acute pyelonephritis in pregnant women often lead to worst health outcome ^1-3^. Chronic pyelonephritis can cause end-stage kidney disease, as often observed in postmenopausal women suffering from a recurrent UTI ^4, 5^. Therefore, it is important to treat UTIs in the early stages of cystitis before the condition progresses to chronic cystitis or pyelonephritis. The innate immunity in the bladder epithelium is crucial as it serves as the first line of defense against uropathogens that can breach this protective barrier to colonize inside the bladder epithelial cells causing UTI ^6, 7^. This bacterial colonization of the bladder epithelium triggers the secretion of both pro-inflammatory and anti-inflammatory cytokines as part of the immediate innate immune responses generated by these epithelial cells ^8-10^. Most UTI cases are caused by uropathogenic *E. coli* (UPEC) that internalize and hide in the bladder epithelial cells using host proteins as their receptors to form intracellular bacterial reservoirs and cause recurrent UTIs ^11, 12^. Dr *E. coli* are among such UPECs that evades the host immune system by internalizing into the uroepithelial cells by binding to CD55 which is a complement regulatory protein expressed on host cell membranes ^13^.

Postmenopausal and pregnant women have been reported to show a higher susceptibility to recurrent UTIs and have the worst disease outcome, suggesting hormonal etiology ^1, 14^. Several clinical and experimental studies have reported the involvement of estrogen in modulating the host immunity in the bladder and dictating the UTI disease outcome ^15-17^. Our previous studies have also shown that estrogen regulates CD55 expression that impacts the Dr *E. coli* colonization ^1, 15^. Estrogen or estrogen receptor (ER)-related mechanisms for modulating the innate immunity in the urinary tract remain poorly understood.

Robust innate immune responses in the bladder are generated by the epithelial cells and resident immune cells that play a major role in protecting the urinary tract from infections ^6, 18^. In response to bacterial infections, resident immune cells and epithelial cells of the bladder release pro-inflammatory cytokines, one of them being TNFα ^19-21^. The local production of TNFα helps transepithelial migration of neutrophils that also assist in the bacterial clearance ^7^. Thus, TNFα is a critical cytokine of the innate immune system as it renders protection against intracellular bacterial infections and also shown to inhibit the development of many inflammatory diseases ^22^. Once the infections are cleared, the epithelial cells and the resident immune cells of the bladder release anti-inflammatory cytokines, like IL10 to reduce the inflammation ^23^. IL10 is another important immunoregulatory cytokine that inhibits activation of macrophages and cytokine production, thus reducing local inflammation ^24, 25^. While higher TNFα expression is necessary for eliminating bacterial infections, induction of IL10 expression after bacterial clearance is required to prevent hyperinflammation that can be induced by uncontrolled TNFα expression. Therefore, maintaining an equilibrium between TNFα and IL10 expression locally as well as systemically is crucial for restoring homeostasis state. Expression of both TNFα and IL10 in tissues and cells of the endometrium, brain, bone and gastrointestinal origin have been found to be differentially regulated by estrogen ^26-28^ via different ERs ^29-35^. Recently, we reported for the first time about the differential involvement of ERα in modulating innate immunity in the bladder versus kidney during Dr *E. coli* induced UTI pathogenesis in an experimental mouse model. ERα played a protective role against infection in the kidney, however, the protection against infection in the bladder was mediated by ER subtypes other than ERα, boosting local immunity via increased expression of TNFα ^36^. However, the involvement of estrogen and ERs in eliciting the TNFα or IL10 expression in the human bladder or kidney epithelium in response to Dr *E. coli* infection has not been studied till date.

Estrogen binds to its receptors, ERα, ERβ or GPR30, to form transcriptional factors that induce the transcription of several estrogen responsive genes involved in immune responses and inflammation ^37^. Differential distribution of these ER subtypes in various mouse and human tissues contribute to variable levels of expression of immune factors in different tissues ^38-42^. ER subtype involvement in modulating the local immunity in the human urinary tract in response to UTI has not been investigated so far. Thus, it is crucial to identify the precise role of each ER subtype in modulating the innate immune responses in the urinary tract that will result in efficient bacterial clearance

Therefore, in the current study we investigated the involvement of all three ER subtypes in regulating the colonization and invasion by Dr *E. coli* in human bladder cells by directly modulating the expression of innate immune cytokines, TNFα and IL10, and bacterial colonization receptor, CD55. This study was conducted in an *in vitro* model established with human bladder epithelial carcinoma (HBEC) 5637 cells. Understanding the ER-regulated innate immune mechanisms operating in the human bladder cells will help us in designing the novel and effective immune based therapies that are highly needed for treating UTIs in women by boosting or regulating the local immune responses in the bladder.

## 2. Materials and methods

### 2.1. Cell culture

The research protocols followed in this study involving the use of human bladder cells and Dr *E. coli* strain were approved by the Institutional Biosafety Committee (IBC) of Oklahoma State University Center for Health Sciences. This study did not require any other ethical board approval because no human subjects or animal tissues were used. Human bladder epithelial carcinoma (HBEC) 5637 cells (kindly provided by Doris M. Benbrook, University of Oklahoma Health Sciences Center) were cultured and maintained in sterile Roswell Park Memorial Institute (RPMI) 1640 medium (Invitrogen, Carlsbad, CA) supplemented with 10% fetal bovine serum (FBS) (Atlanta Biologicals, Flowery Branch, GA), 100 U/ml penicillin, 100 µg/ml streptomycin (Invitrogen, Carlsbad, CA), 1 mM sodium pyruvate (Invitrogen, Carlsbad, CA) and 1% GlutaMAX™ Supplement (Invitrogen, Carlsbad, CA) incubated at 37°C in a humidified atmosphere (95%) containing 5% CO_2_.

### 2.2. Hormonal drug treatment of bladder cells

HBEC 5637 cells were plated at different densities in 6-well, 12-well, 24-well plates and 96-well plates for various assays and were allowed to grow overnight (18–24 hours). Before hormone drug treatment, the cells were subjected to serum starvation for 20-24 hours. After serum starvation, the cells were incubated with different doses of various drugs at for another 24 hours till further experimentation. The hormonal drugs that were used include, Propylpyrazole Triol (PPT), Methyl-piperidino-pyrazole (MPP), 2,3-*bis* (4-Hydroxyphenyl) propionitrile (DPN), 4-[2-Phenyl-5,7-bis(trifluoromethyl)pyrazolo[1,5-a]pyrimidin-3-yl]pheno) (PHTPP), (±)-1-[(3aR*,4S*,9bS*)-4-(6-Bromo-1,3-benzodioxol-5-yl)-3a,4,5,9b-tetrahydro-3H-cyclopenta[c]quinolin-8-yl]-ethanone (G1) and (3aS*,4R*,9bR*)-4-(6-Bromo-1,3-benzodioxol-5-yl)-3a,4,5,9b-3H-cyclopenta[c]quinoline (G15). These drugs were purchased from Cayman Chemicals (Ann Arbor, MI). The stock solutions were prepared in DMSO and diluted to the required working concentrations in serum-free media. The vehicle control (C) included in the assay had only serum free culture medium with DMSO (0.01% vol/vol).

### 2.3. Gene-silencing by siRNA transfection

HBEC 5637 cells were grown to 60% confluency before transfection with either *esr1*-targeting siRNAs (ID-s4825, Cat. no. 4392420, Ambion, Waltham, MA) or non-specific (negative control or NC) siRNA (Cat. no. 4390843, Ambion, Waltham, MA) using lipofectamine reagent (Invitrogen, Waltham, MA) following the manufacturer’s instructions. After 48 hours, *esr1* gene silencing was confirmed by both quantitative real-time RT-PCR and immunofluorescence staining. These *esr1*^-/-^ bladder cells were subjected to further experimentation.

### 2.4. Cell viability assay

HBEC 5637 cells were grown overnight in 96-well plates (50,000 cells/ well) and treated with different concentrations of either hormonal drugs for 24 hours or siRNA transfection reagent for 48 hours. The cells were then treated with MTT solution (0.5 mg/ml in serum free RPMI media) for 2 hours at 37°C. The formazan compound formed in the cells after 2 hours was solubilized in a 3:1 mixture of DMSO and serum free RPMI media and the OD was measured at 540 nm. The non-toxic drug doses that were selected for the treatment of cells in experimental assays were as follows: PPT (0.01 µM, 0.1 µM, and 1 µM), MPP (0.01 µM, 0.1 µM, and 1 µM), DPN (0.01 µM, 0.1 µM, 1 µM and 10 µM), PHTPP (0.001 µM, 0.01 µM, 0.05 µM and 0.1 µM), G1 (0.001 µM, 0.01 µM, 0.1 µM) and G15 (0.1 µM, 1 µM, 2.5 µM). The optimum non-toxic concentration of *esr1* siRNA and negative control siRNA used for experimental assays was 10 nM.

### 2.5. Preparation of bacterial inoculum

The presence of Dr adhesin on Dr *E. coli* strain-IH11128 (O75:K5: H^-^ strain) was confirmed by hemagglutination of human O group erythrocytes as described previously ^13^. Isolated colonies of Dr *E. coli* and its isogenic mutant Dr ^-^ *E. coli* from overnight cultures were resuspended in serum free RPMI media. A suspension of Dr *E. coli* with an OD of 0.5 measured at 600 nm (4.5 × 10^8^ cfu/ml) was prepared for the infection of bladder cells.

### 2.6. Determination of bacterial invasion by Gentamicin Protection Assay

HBEC 5637 cells were grown overnight in 24-well plates (150,000 cells/well). Upon reaching 70% confluency, the cells were subjected to either 24-hour drug treatment or 48-hour of siRNA transfection. The cells were then infected with Dr *E. coli* suspension at a multiplicity of infection (MOI) of approximately 30. The cell culture plates were centrifuged at 500 x g and incubated for 1 hour at 37°C. Bacterial suspension was removed and the cells were washed once with sterile PBS. To kill extracellular bacteria, the cells were incubated with 100 µg/ml gentamicin for 1 hour. Gentamicin was aspirated and the cells were washed with sterile PBS to remove any residual gentamicin. The cells were then lysed with buffer containing 1% Triton X-100 in PBS. The cell lysates were plated on LB agar plates and incubated at 37°C overnight to determine the number of viable bacteria internalized in the cells. Bacterial colonies were counted and represented as colony-forming units (CFU). The results of gentamicin protection assay were reported as percentage of bacterial invasion in treated cells relative to bacterial invasion in untreated control (C) cells (considered as 100%). The gentamicin protection assay for each treatment condition was performed 6-8 times with 3 replicates per experiment.

### 2.7. RNA isolation, cDNA synthesis and quantitative RT-PCR analyses

Total RNA was isolated from drug treated or siRNA transfected HBEC 5637 cells using TRIzol reagent (Life Technologies, Grand Island, NY) and cDNA was synthesized. Quantitative RT-PCR was performed using PowerUp™ SYBR® Green Master Mix (Applied Biosystems, Foster City, CA). The expression levels of target genes, *cd55, tnfa* and *il10* were normalized to the endogenous control gene, *gapdh* and to the experimental control (C) and reported as 2^-ΔΔCt^ values. Quantitative RT-PCR was carried out using 7500 Real-Time PCR System (Applied Biosystems, Foster City, CA). The primer pairs were purchased from Integrated DNA Technologies (Coralville, IA) and are listed below:

*cd55* (Forward primer - 5’ TTTCCAGGACAACCAAGCATT 3’, Reverse primer - 5’ ACACGTGTGCCCAGATAGA 3’),

*tnfa* (Forward primer-5’ TGTAGCCCATGTTGTAGCAAAC 3’, Reverse primer-5’ AGAGGACCTGGGAGTAGATGA 3’)

*il10* (Forward primer-5’ AATGAAGGATCAGCTGGACAAC 3’, Reverse primer-5’ CCAGGTAAAACTGGATCATCTCAG 3’)

*gapdh* (Forward primer-5’ GCACCGTCAAGCTGA 3’, Reverse primer-5’ ACTCAGCGCCAGCATC 3’),

*esr1* (Forward primer-5’ CCAACCAGTGCACCATTGAT 3’, Reverse primer-5’ GGTCTTTTCGTATCCCACCTTT 3’),

*esr2* (Forward primer-5’ GGCAGAGGACAGTAAAAGCA3’, Reverse primer-5’ GGACCACACAGCAGAAAGAT3’),

*gper1* (Forward primer-5’ GTACTTCATCAACCTGGCGGTG3’, Reverse primer-5’ TCATCCAGGTGAGGAAGAAGACG3’)

### 2.8. Immunofluorescence staining of cells for epifluorescent microscopy

HBEC 5637 cells were grown overnight on glass coverslips (#1.5) placed inside in each well of 12-well plate (150,000 cells/well) incubated overnight at 37 °C with 5% CO_2_. The cells were then subjected to either 24-hour drug treatment or 48-hour of siRNA transfection. Upon reaching 70-75% confluency, the cells were rinsed 3 times with ice-cold PBS and fixed in 4% paraformaldehyde for 1 hour at room temperature. Only the cells to be stained for nuclear proteins were permeabilized with 0.1% Triton X-100 for 10 minutes at room temperature. Both permeabilized and non-permeabilized cells were then incubated in 3% BSA for 30 minutes at room temperature to block non-specific binding sites. After blocking, the cells were incubated with primary antibodies diluted in 3% BSA and left overnight at 4°C. The primary antibodies used were rabbit anti-human ERα (ab108398) at 1:50 dilution, rabbit anti-human ERβ (ab3577) at 1:2000 dilution, rabbit anti-human GPR30 (ab39742) at 1:400 dilution and rabbit anti-human CD55 (ab1422) at 1:200 dilution. Next day, the cells were washed three times with PBS and further incubated with FITC labelled goat anti-rabbit IgG (ab 150077) prepared in 3% BSA at a dilution of 1:500 left for 1 hour in the dark at room temperature. After a washing step, the coverslips were mounted on a glass slide with Prolong Diamond Antifaded mounting reagent (Thermofisher, Waltham, MA) containing DAPI to counterstain nuclei. Samples were imaged using a Nikon Eclipse Ts2R epifluorescent microscope.

### 2.9. Western Blotting Assays

Drug treated or siRNA transfected HBEC 5637 cells were collected from 6-well plates (containing 300,000 cells/well) using cell dissociation buffer (Life Technologies, Waltham, MA). The cells were lysed with RIPA lysis buffer supplemented with Halt™ Protease and Phosphatase Inhibitor Cocktail (Thermofisher, Waltham, MA) and phenylmethylesulfonyl fluoride (Thermofisher, Waltham, Ma). The total protein concentration in each cell lysate was measured using Pierce BCA Protein Assay Kit (Thermofisher, Waltham, MA). The cell lysates containing approximately 20 µg of protein were resolved on Novex™ NuPAGE™ 4-12% Bis-Tris Protein Gels (Invitrogen, Waltham, MA). The separated proteins were transferred to nitrocellulose membranes that were subsequently blocked using 5% non-fat dry milk diluted in TBST (10 mM Tris-HCL, 100 mM NaCl, 0.2% Tween-20) at room temperature for 2 hours. After blocking, the membranes were incubated with rabbit anti-human CD55 antibody (ab133684) at a dilution of 1:5000 and rabbit anti-human β-actin antibody (ab8227) at a dilution of 1:1000 overnight at 4°C. Both antibodies were diluted in TBST containing 5% milk. Membranes were subsequently probed with an alkaline phosphatase conjugated goat anti-rabbit IgG (7054S, Cell Signaling) used at a dilution of 1:1000 in TBST containing 5% milk at room temperature for 2 hours. The immunoreactivity was carried out using an Enhanced Chemiluminescence reagent (Fisher Scientific, Hampton, NH).

### 2.10. Statistics

GraphPad Prism 7.03 (Graph Pad software Inc.) was used for statistical comparisons between experimental groups. Statistical analysis for more than two experimental groups was conducted using parametric one-way ANOVA followed by Dunnet’s post-hoc test, for multiple comparisons. A p-value of less than 0.05 was considered significant.

## 3. Results

### 3.1. HBEC 5637 cells express CD55 which serves as a binding ligand for Dr E. coli during cell invasion and infection

Before establishing the *in vitro* model of UTI pathogenesis using HBEC 5637 cells, we confirmed the expression of CD55 in these cells as CD55 serves as colonization receptor for Dr *E. coli* ^13, 43^. We observed a high expression of CD55 protein in these cells (Figure 1 a). In order to confirm the colonization and invasion by Dr *E. coli* in these cells is mediated by binding to CD55, we blocked CD55 expression in these cells by pre-treatment of cells with CD55 antibody prior to bacterial invasion. Dr *E. coli* invasion in cells was significantly reduced in a dose-dependent manner by antibody dilutions of 1:200 (P < 0.01), 1:100 (P < 0.001), 1:50 and 1:20 (P < 0.0001) (Figure 1 b), confirming that CD55 serves as the colonization receptor for Dr *E. coli* in 5637 cells. To further confirm that bacterial invasion in HBEC cells was facilitated by the presence of Dr fimbriae, we included Dr^**-**^ *E. coli* (an isogenic mutant for Dr fimbriae) strain as a negative control in the bacterial invasion assay. Dr *E. coli* invasion in cells pre-treated with CD55 antibody was found to be significantly low (P < 0.0001) when compared to antibody untreated Dr *E. coli* infected cells (Figure 1 c), confirming that bacterial invasion in HBEC 5637 cells is mediated via CD55. Further, a significantly low (P < 0.0001) Dr^**-**^ *E. coli* invasion was observed in antibody untreated cells and in cells pre-treated with CD55 antibody, when compared to untreated Dr *E. coli* infected cells (Figure 1 c). This observation further confirms that the presence of both Dr fimbriae and CD55 are crucial for Dr *E. coli* invasion in HBEC 5637 cells.

**Figure 1:**
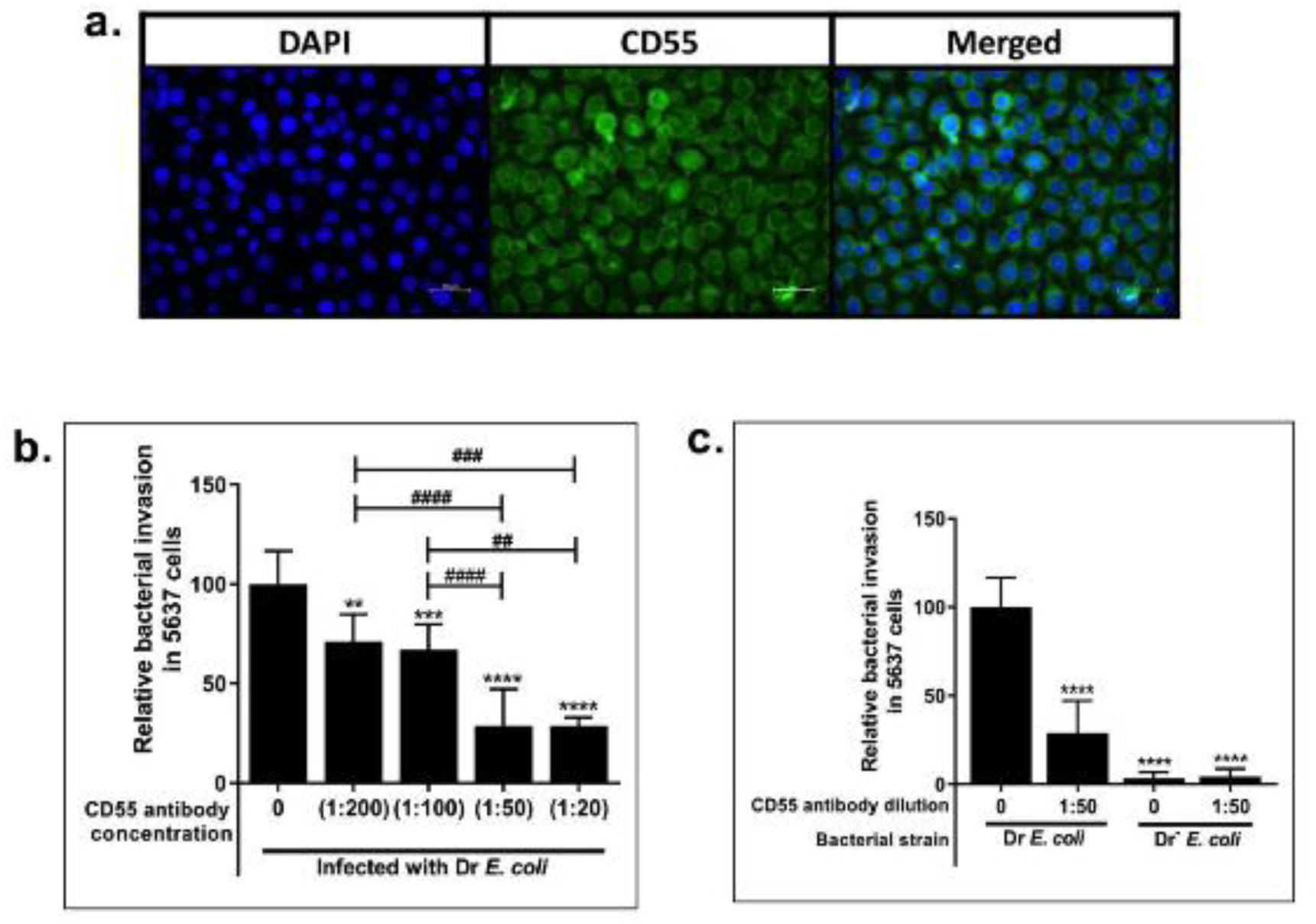
**(a) Expression of CD55 in HBEC 5637 cells detected by immunofluorescence staining. (b) Relative invasion by Dr *E. coli* in cells pre-incubated with CD55 antibody for blocking CD55 receptor.** Cells incubated with CD55 antibody had significantly low bacterial invasion. **(c) Relative invasion by Dr *E. coli* and Dr^−^ *E. coli* in HBEC 5637 cells with or without CD55 antibody pre-treatment.** Dr^−^ *E. coli* showed significantly low bacterial invasion as compared to Dr *E. coli. (**P<0.01, ***P<0.001, ****P<0.0001 indicate significant differences between antibody treated group and the control group*. ^*# #*^*P<0.01*, ^*# # #*^*P<0.001*, ^*# # # #*^*P<0.0001, indicate significant differences between antibody treated groups as indicated)*

### 3.2. HBEC 5637 cells express all three ER subtypes, ERα, ERβ and GPR30

The mRNA and protein expressions of the three ER subtypes in HBEC 5637 cells was assessed. The expression of *esr2* mRNA was 1.5-fold higher than expression of *esr1* mRNA (P < 0.001; Figure 2 a). ERβ protein expression was 2-fold higher than ERα (P < 0.0001; Figure 2 b). Similarly, *gper1* mRNA expression was also 2 fold higher than *esr1* mRNA (P < 0.001; Figure 2 a). GPR30 protein expression in cells was 2.5 fold higher than ERα expression (P < 0.001; Figure 2 b). A significantly higher mRNA (P < 0.05) and protein (P < 0.0001) expression of GPR30 was also observed in these cells when compared to ERβ (Figures 2 a and 2 b).

**Figure 2:**
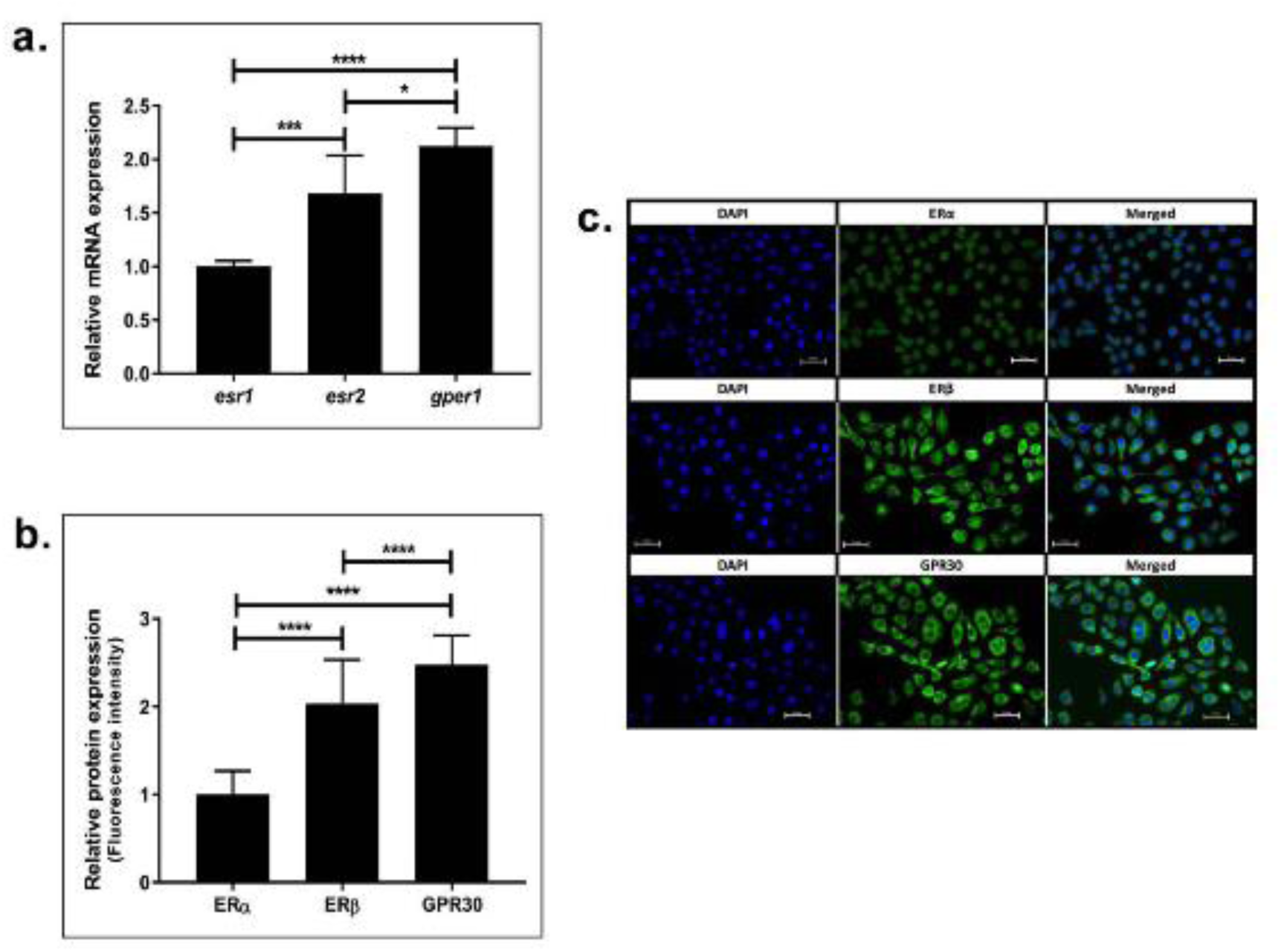
Expression of estrogen receptor subtypes in HBEC 5637 cells. (a) *esr2* and *gper* mRNA expression were significantly higher compared to *esr1* mRNA expression levels. (b) ERβ and GRP30 protein levels were also significantly higher than ERα. (c) Representative images showing the protein expression of estrogen receptors detected by immunofluorescence staining. (*\*P<0.05, ***P<0.001, ****P<0.0001 indicate significant differences between ERα, ERβ and GRP30 expression)*

### 3.3. Drug-induced modulation of ER subtype activity in HBEC 5637 cells differentially regulate Dr E. coli invasion

We determined the role of each ER subtype in modulating the Dr *E. coli* invasion in HBEC 5637 cells pre-treated with specific ER subtype agonists or antagonists. ERα agonist, PPT treatment of cells led to a dose-dependent significant inhibition of bacterial invasion (70-85%) observed at all three doses, 0.01 µM, 0.1 µM and 1 µM (P < 0.0001) compared to vehicle treated control (Figure 3 a). ERα antagonist, MPP treatment reversed the protection mediated by PPT in a dose-dependent manner. However, MPP treatment of cells resulted in a partial protection with only 40-60% reduction in bacterial invasion at all three doses, 0.01 µM, 0.1 µM and 1 µM when compared to control (P < 0.0001) (Figure 3 a).

**Figure 3:**
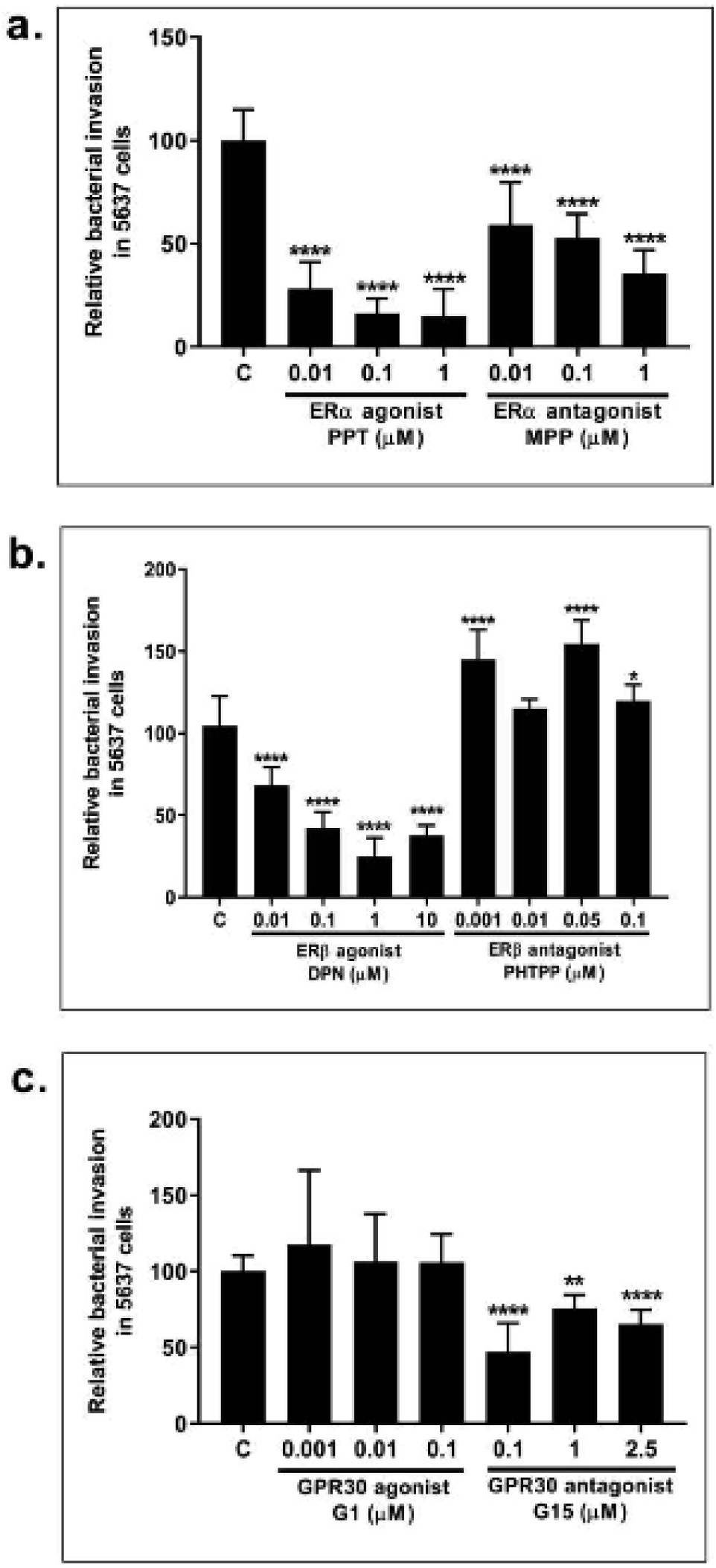
Relative invasion by Dr *E. coli* in HBEC 5637 cells pre-treated with (a) ERα agonist PPT and ERα antagonist MPP, (b) ERβ agonist DPN and ERβ antagonist PHTPP, (c) GPR30 agonist G1 and GPR30 antagonist G15. PPT was most efficient in inhibiting bacterial invasion followed by DPN and G15. PHTPP was least effective in inhibiting bacterial invasion followed by G1 and MPP. (*\*P<0.05, ***P<0.001, ****P<0.0001 indicate significant differences between drug treated groups and vehicle control (C) group)*

ERβ agonist, DPN treatment of cells also resulted in a significant reduction in bacterial invasion ranging between 30-70% at all doses (P < 0.0001), with 1 µM DPN being the most protective resulting in 70% reduction in bacterial invasion. In contrast, ERβ antagonist, PHTPP treatment reversed the protection mediated by DPN. PHTPP led to a significant increase in bacterial invasion ranging between 20-55% at doses 0.001 µM (P < 0.0001), 0.05 µM (P < 0.0001) and 0.1 µM when compared to control (P < 0.05), except at 0.01 µM dose, which was comparable to control.

In contrast, the activation of GPR30 by agonist G1 treatment of cells did not result in any protection as observed with ERα and ERβ activation. Increased bacterial invasion ranging between 10-20% was observed in cells at all the doses of G1, 0.001 µM, 0.1 µM and 1 µM when compared to control (P > 0.05). However, blocking of GPR30 in cells by antagonist G15 resulted in a significantly low bacterial invasion ranging between 20-50% at all doses, 0.1 µM (P < 0.0001), 1 µM (P < 0.01) and 2.5 µM (P < 0.0001).

In summary, ERα agonist, PPT was the most efficient in protecting against bacterial invasion in bladder cells, followed by ERβ agonist, DPN and GPR30 antagonist, G15. The least protection was mediated by suppression of ERβ by antagonist, PHTPP, followed by activation of GPR30 by agonist, G1 and suppression of ERα by its antagonist, MPP.

### 3.4. Drug-induced modulation of ER subtype activity in HBEC 5637 cells differentially regulates CD55 expression

In our previous experiments, treatment of cells with specific ER subtype agonists or antagonists led to differential protection against bacterial invasion. Hence, we further investigated the effects of these drugs on CD55 mRNA and protein expression as CD55 facilitates Dr *E. coli* colonization in cells.

ERα agonist, PPT treatment significantly reduced *cd55* mRNA expression in HBEC 5637 cells at doses 0.01 µM, 0.1 µM and 1 µM (P < 0.0001) (Figure 4 a) when compared to control. ERα antagonist, MPP treatment also significantly reduced *cd55* mRNA expression in these cells at doses 0.01 µM, 0.1 µM and 1 µM (P < 0.0001) (Figure 4 a), however, not to the extent observed in PPT treated cells, explaining the partial reversal of protection.

**Figure 4:**
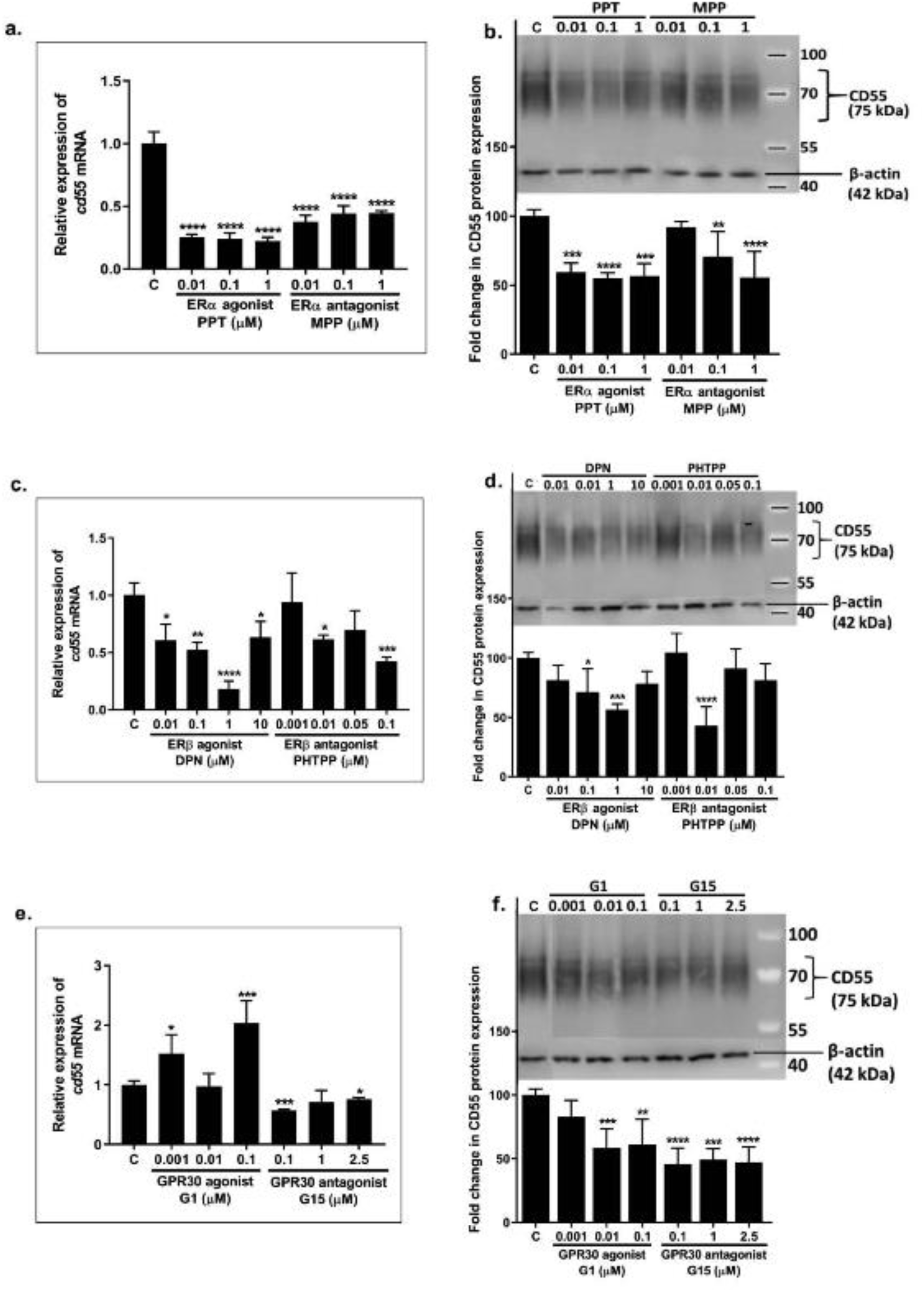
Relative expression of *cd55* mRNA and CD55 protein in HBEC 5637 cells pre-treated with (a and b), ERα agonist PPT and ERα antagonist MPP, (c and d) ERβ agonist DPN and ERβ antagonist PHTPP, (e and f) GPR30 agonist G1 and GPR30 antagonist G15. CD55 mRNA or CD55 protein expression corresponded to bacterial invasion at each drug dose. (*\*P<0.05, **P<0.01, ***P<0.001, ****P<0.0001 indicate significant differences between drug treated groups and vehicle control (C) group)*

CD55 protein expression was significantly reduced in PPT treated cells at doses 0.01 µM (P < 0.001), 0.1 µM (P < 0.0001) and 1 µM when compared to control, (P < 0.001) (Figure 4 b). In contrast, MPP treatment of cells increased CD55 protein expression when compared to CD55 expression by PPT, explaining the reversal of protection. However, when compared to control, MPP treatment significantly reduced CD55 protein expression compared to control at 0.1 µM (P < 0.01) and 1 µM (P < 0.0001) (Figure 4 b).

ERβ agonist, DPN treatment of cells significantly reduced *cd55* mRNA expression at doses 0.01 µM (P < 0.05), 0.1 µM (P < 0.01), 1 µM (P < 0.0001) and 1 µM (P < 0.05) when compared to control (Figure 4 c). Expression of *cd55* mRNA in response to ERβ antagonist, PHTPP treatment of cells was higher than 1 µM DPN but comparable to control at dose 0.001 µM and 0.05 µM (P > 0.05). Further, PHTPP treatment significantly reduced *cd55* mRNA expression in cells at doses 0.01 µM (P < 0.05), and 0.1 µM (P < 0.001) (Figure 4 c).

DPN treatment of cells reduced CD55 protein expression at all the doses, however a significant reduction was only observed at 0.01 µM (P < 0.05) and 1 µM (P < 0.001) (Figure 4 d). In contrast, CD55 protein expression in ERβ antagonist, PHTPP treated cells was found to be comparable to control at all doses except for 0.01 µM, which significantly reduced (P < 0.0001) the CD55 expression (Figure 4 d). This significant reduction in CD55 protein expression in response to 0.01 µM PHTPP treatment in cells corresponded with the decrease in bacterial invasion observed at this dose. Changes in both CD55 mRNA and protein expression in response to different doses of DPN and PHTPP were found to be directly correlating with the changes in bacterial invasion at the same doses.

GPR30 agonist, G1 treatment of cells significantly increased *cd55* mRNA expression at doses 0.001 µM (P < 0.05) and 0.01 µM (P < 0.001) when compared to control (Figure 4 e). However, *cd55* mRNA expression levels in response to 0.01 µM G1 treatment was similar to control. In contrast, GPR30 antagonist, G15 treatment reduced *cd55* mRNA expression at all doses, but significantly only at 0.1 µM (P < 0.001) and 2.5 µM (P < 0.05) (Figure 4 e).

Further, G1 treatment of cells reduced CD55 protein expression at all doses as compared to control but significant reduction was only observed at doses 0.01 µM (P < 0.001) and 0.1 µM (P < 0.01) (Figure 4 f). However, GPR30 antagonist, G15 treatment of cells significantly reduced CD55 protein expression at all three doses, 0.1 µM (P < 0.0001), 1 µM (P < 0.001) and 2.5 µM (P < 0.0001) (Figure 4 f). CD55 protein expression in G15 treated cells was lower than that observed in G1 treated cells, explaining the observed protection against bacterial invasion mediated by G15, but not by G1.

### 3.5. Modulation of ER subtype activity in HBEC 5637 cells differentially regulates the gene expression of cytokines tnfa and il10

Pro-inflammatory cytokine, TNFα, plays a major role in eliminating bacterial infections by inducing innate immunity resulting in local inflammation and neutrophil recruitment. In contrast, anti-inflammatory cytokine, IL10 suppresses local inflammation, thus restoring homeostasis after bacterial infections are cleared. In our previous experiments, we observed differential protection against bacterial invasion that was induced by specific ER subtype agonists and antagonists. Hence, we investigated the effects of ER subtype specific agonists and antagonists on TNFα and IL10 cytokine gene expression in bladder cells before and after bacterial infection.

In uninfected HBEC 5637 cells, PPT treatment of cells induced a significant increase in *tnfa* mRNA expression at doses 0.01 µM (P < 0.0001), 0.1 µM (P < 0.0001) and 1 µM (P < 0.001) (Figure 5 a). However, PPT resulted in a significant decrease of *il10* mRNA expression at all doses (P < 0.0001) (Figure 5 a). In contrast, MPP treatment in uninfected cells significantly inhibited *tnfa* mRNA expression at doses 0.01 µM (P < 0.01), 0.1 µM (P < 0.05) and 1 µM (P < 0.001) but significantly increased (P < 0.001) *il10* mRNA expression at all doses when compared to PPT treated group, explaining the reversal of PPT-mediated protection against bacterial invasion (Figure 5 a).

**Figure 5:**
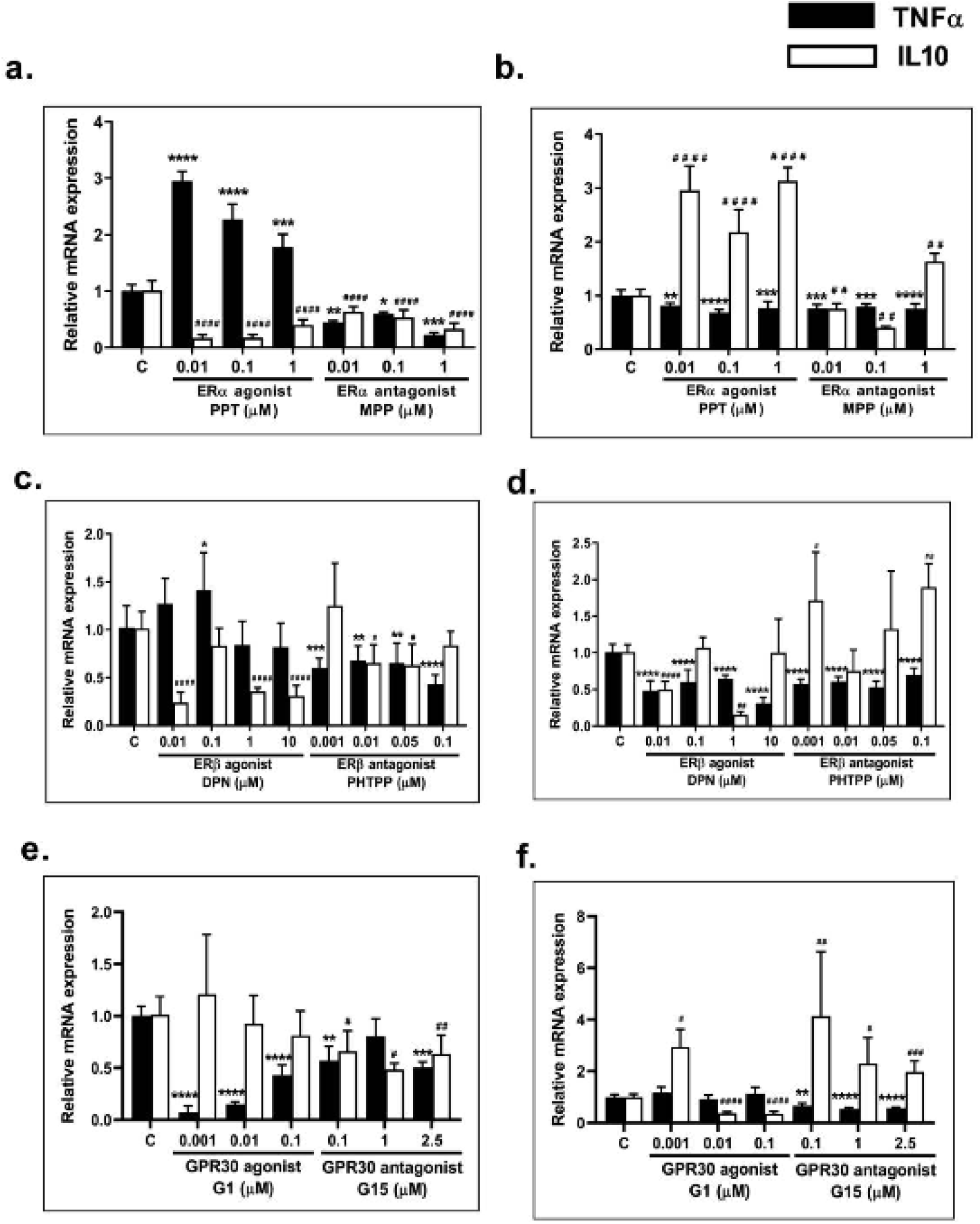
Relative expression of *tnfa* and *il10* mRNA before and after infection in HBEC 5637 cells pre-treated with (a and b), ERα agonist PPT and ERα antagonist MPP, (c and d) ERβ agonist DPN and ERβ antagonist PHTPP, (e and f) GPR30 agonist G1 and GPR30 antagonist G15. *tnfa* mRNA expression before infection corresponded to bacterial invasion at each drug dose. *(*P<0.05, **P<0.01, ***P<0.001, ****P<0.0001, indicate significant differences between tnfa mRNA expression in drug treated groups and vehicle control (C) group*. ^*#*^*P<0.05*, ^*# #*^*P<0.01*, ^*# # #*^*P<0.001*, ^*# # # #*^*P<0.0001, indicate significant differences between il10 mRNA expression in drug treated groups and vehicle control (C) group)*.

Post-infection, the *tnfa* mRNA expression in PPT treated cells reduced significantly, when compared to control at all doses, 0.01 µM (P < 0.05), 0.1 µM (P < 0.0001) and 1 µM (P < 0.001) (Figure 5 b). However, the *il10* mRNA expression in these cells significantly increased (P < 0.0001) at all doses (Figure 5 b). In contrast, *tnfa* mRNA expression in MPP treated cells increased after infection but were still significantly lower than the control at all doses, 0.01 µM (P < 0.001), 0.1 µM (P < 0.001) and 1 µM (P < 0.0001) (Figure 5 b). Post-infection, the *il10* mRNA expression was significantly reduced (P < 0.01) in MPP treated cells (Figure 5 b), when compared to control and PPT treated cells explaining the higher bacterial invasion observed in MPP treated cells.

In uninfected cells, ERβ agonist, DPN treatment significantly increased *tnfa* mRNA expression only at doses 0.1 µM (P < 0.05) and 0.01 µM, but the *tnfa* mRNA expression at doses 1 µM or 10 µM were comparable to control (Figure 5 c). However, DPN treatment in uninfected cells significantly reduced (P < 0.0001) *il10* mRNA expression at doses 0.01 µM, 1 µM and 10 µM, but the expression at 0.1 µM was comparable to control (Figure 5 c). In contrast, ERβ antagonist, PHTPP treatment of cells significantly reduced *tnfa* mRNA expression in cells at all doses 0.001 µM (P < 0.001), 0.01 µM (P < 0.01), 0.05 µM (P < 0.01) and 0.1 µM (P < 0.0001) when compared to control (Figure 5 c). However, PHTPP treatment of cells increased *il10* mRNA expression only at 0.001 µM, but significantly reduced (P < 0.05) *il10* mRNA expression at 0.01 µM and 0.05 µM (Figure 5 c). The *il10* mRNA expression in response to 0.1 µM PHTPP was comparable to control. Further, *tnfa* mRNA expression levels observed in response to different doses of PHTPP treatment were still lower than the expression levels observed with DPN treatment. Also, *il10* mRNA expression levels observed in response to PHTPP treatment of cells at different doses were higher than the expression levels observed with DPN treatment. These opposite effects on *tnfa* and *il10* mRNA expression in DPN versus PHTPP treated bladder cells explain the protective effects of DPN against bacterial invasion when compared to PHTPP treated cells.

Post-infection, the *tnfa* mRNA expression levels were significantly lower in both DPN and PHTPP treated cells at all doses (P < 0.0001) compared to control (Figure 5 d). However, *tnfa* mRNA expression levels observed in DPN treated infected cells were comparable to the levels observed in PHTPP treated infected cells. The *il10* mRNA expression in DPN treated infected cells was significantly reduced at doses 0.01 µM (P < 0.0001) and 1 µM (P < 0.01), but the expression levels at 0.1 µM and 10 µM DPN treatment were comparable to control. In contrast, *il10* mRNA expression levels in PHTPP treated infected cells were higher than control at doses 0.001 µM (P < 0.05), 0.05 µM and 0.1 µM (P < 0.01), but comparable to control at 0.01 µM. Also, the *il10* mRNA expression levels in DPN treated infected cells were lower than the expression levels in PHTPP treated infected cells.

In uninfected HBEC 5637 cells, GPR30 agonist, G1 treatment led to a significantly low *tnfa* mRNA expression at all doses (P < 0.0001) when compared to control (Figure 5 e). No changes in *il10* mRNA expression were observed in response to G1 treatment in these cells (Figure 5 e). In contrast, GPR30 antagonist, G15 treatment of cells increased *tnfa* mRNA expression levels and reduced *il10* mRNA expression levels when compared to G1 treatment (Figure 5 e).

Post-infection, *tnfa* mRNA expression in cells were minimally higher in G1 treated groups when compared to control (Figure 5 f). In contrast, the *il10* mRNA expression in these cells significantly reduced with G1 treatment at doses 0.01 µM and 0.1 µM (P < 0.0001) (Figure 5 f). However, *tnfa* mRNA expression levels significantly reduced in G15 treated infected cells at doses 0.1 µM (P < 0.01), 1 µM and 2.5 µM when compared to control (P < 0.0001), but the *il10* mRNA expression significantly increased in these cells at doses 0.1 µM (P < 0.01), 1 µM (P < 0.05) and 2.5 µM (P < 0.001) when compared to control (Figure 5 f). These opposite effects of G1 or G15 treatment on *tnfa* and *il10* mRNA expression in infected cells corresponded with the bacterial invasion outcome that was observed in response to treatment with these drugs.

### 3.6. siRNA-mediated targeting of esr1 mRNA in HBEC 5637 cells leads to suppression of ERα expression

Our results from the Dr *E. coli* bacterial invasion studies in HBEC 5637 cells showed that ERα agonist was the most protective against the bacterial invasion followed by ERβ agonist and GPR30 antagonist. A complete reversal of this protection against bacterial invasion was observed only in response to treatment with ERβ antagonist and GPR30 agonist. However, ERα antagonist, MPP only showed partial reversal of protection mediated by ERα agonist, PPT against bacterial invasion. Hence, to further confirm these results related to ERα, direct silencing of *esr1* gene expression using siRNA was undertaken in 5637 bladder cells before infection with Dr *E. coli*.

Our results showed a significant decrease (P < 0.0001) in *esr1* mRNA (Figure 6 a) expression levels and ERα protein (Figure 6 b and c) expression in cells treated with siRNA as compared to no treatment control or negative control (treated with non-specific RNA).

**Figure 6:**
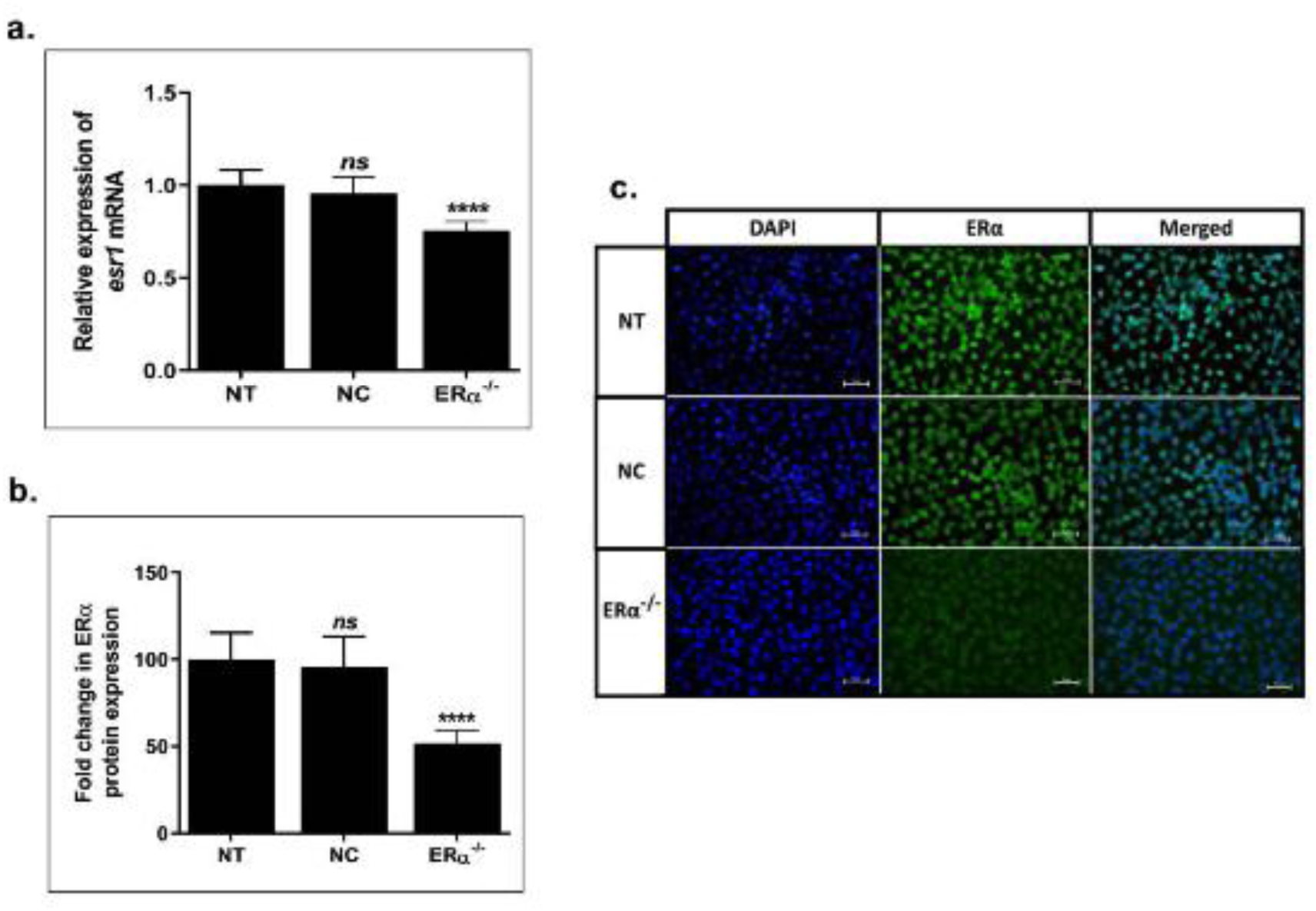
Silencing of ERα protein in HBEC 5637 cells using siRNA. Expression of (a) *esr1* mRNA and (b) ERα protein significantly reduced in cells treated with ERα siRNA as compared to no treatment (NT) control and negative control (NC) cells treated with non-specific siRNA. (c) Representative images showing reduction of ERα protein detected by immunofluorescence staining. (*\*\*\*\*P<0.0001, indicate significant differences relative to no treatment control (NT), ns-not significant*.

### 3.7. siRNA-induced silencing of esr1 gene expression in HBEC 5637 cells lead to increased bacterial invasion and CD55 expression

We determined the effects of *esr1* gene silencing on Dr *E. coli* invasion in HBEC 5637 cells. Silencing of *esr1* gene in these cells significantly increased (P < 0.0001) Dr *E. coli* invasion by 180% when compared to both the controls (Figure 7 a), indicating the protective role of ERα against Dr *E. coli* invasion. Silencing of *esr1* in cells did not affect the expression of *cd55* mRNA (Figure 7 b), however CD55 protein expression was found to be significantly increased (P < 0.05) in *esr1*^-/-^ 5637 cells when compared to both control groups (Figure 7 c). This further explains the increased bacterial invasion that was observed in *esr1*^*-/-*^ cells.

**Figure 7:**
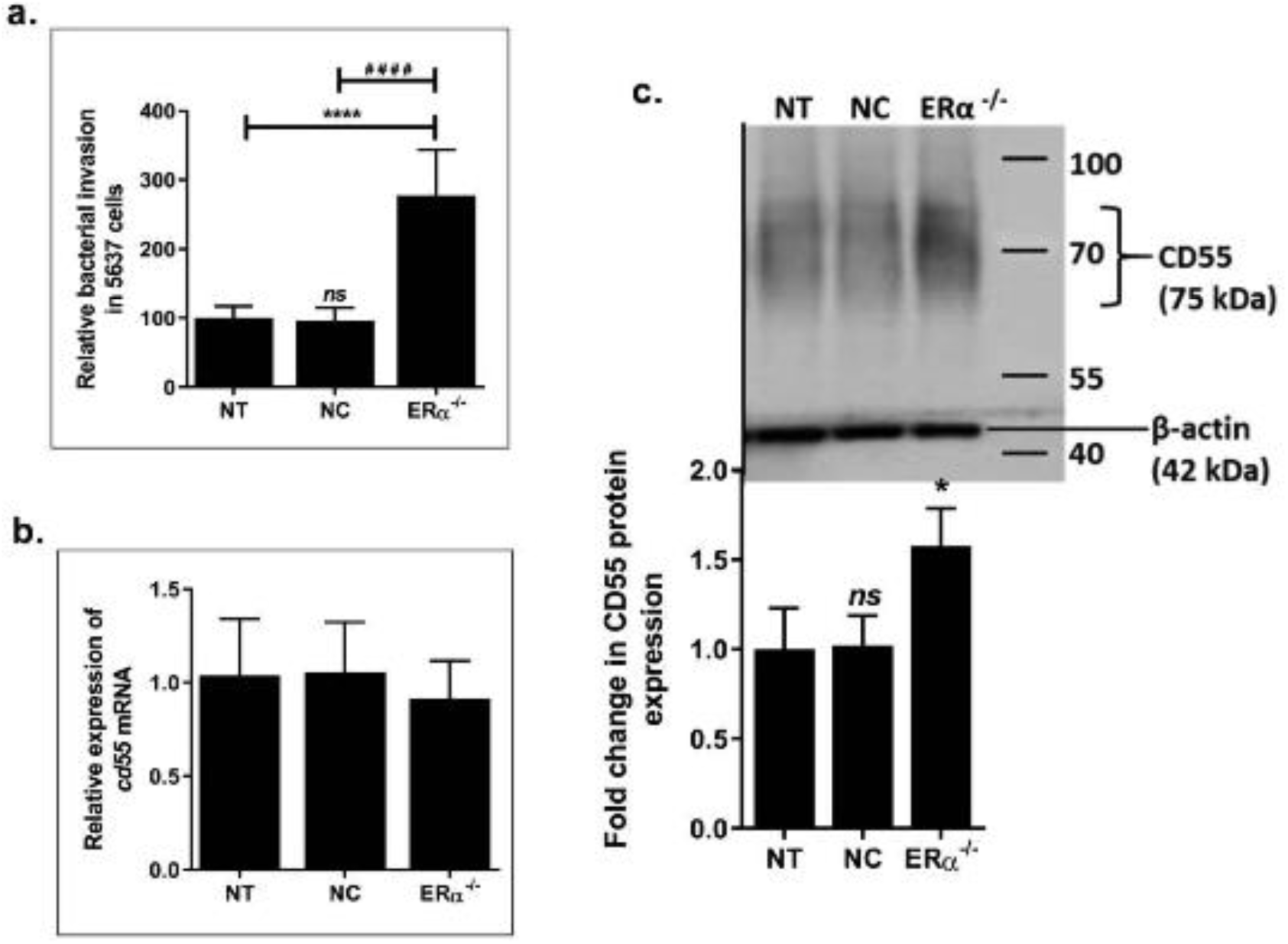
Relative invasion by Dr *E. coli* (a) and relative expression of CD55 (b) mRNA and (c) protein in ERα^-/-^ HBEC 5637 cells. Bacterial invasion and CD55 mRNA and protein expression in ERα^-/-^ cells were significantly higher as compared to both controls. (*\*P<0.05, ****P<0.0001, indicate significant differences relative to no treatment control (NT), ns-not significant)*.

### 3.8. siRNA-induced silencing of ERα in HBEC 5637 cells differentially regulates the gene expression of cytokines tnfa and il10

The *tnfa* and *il10* mRNA expression levels in *esr1*^*-/-*^ cells were determine before (Figure 8 a) and after (Figure 8 b) infection. In uninfected state, *tnfa* mRNA expression in *esr1*^*-/-*^ cells were significantly suppressed (P < 0.0001) and the *il10* mRNA expression levels were significantly elevated (P < 0.05) when compared to controls. Low *tnfa* mRNA expression levels and high *il10* mRNA expression levels in *esr1*^*-/-*^ cells explain the observed high Dr *E. coli* invasion in these cells. The *tnfa* mRNA expression levels in infected *esr1*^*-/-*^ cells were significantly suppressed (P < 0.01) but not to the extent as observed in uninfected *esr1*^*-/-*^ cells. However, the *il10* mRNA expression levels were found to be significantly low (P < 0.01) when compared to controls.

**Figure 8:**
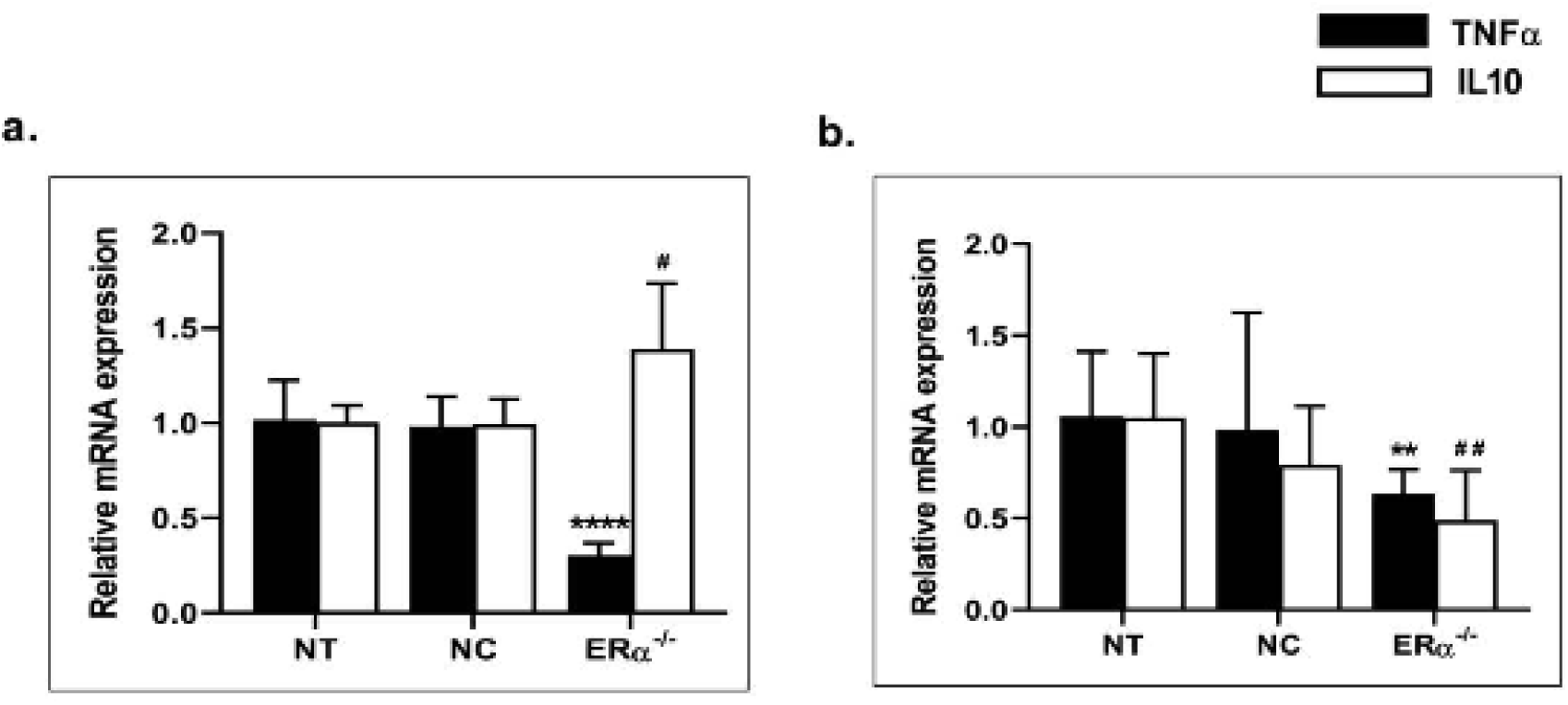
Relative expression of *tnfa* and *il10* mRNA in ERα^-/-^ HBEC 5637 cells (a) before and (b) after infection. Expression of *tnfa* and *il10* mRNA in ERα^-/-^ cells were significantly lower as compared to both controls. *(**P<0.01, ****P<0.0001, indicate significant differences between tnfa mRNA expression in siRNA treated groups and no treatment control (NT) group*. ^*#*^*P<0.05*, ^*# #*^*P<0.01, indicate significant differences between il10 mRNA expression in siRNA treated groups and no treatment control (NT) group)*.

## 4. Discussion

The development of early innate immune responses in the bladder epithelium serves as the first line of defense against colonization and invasion by uropathogens thus dictating the UTI disease outcome. Some of the uropathogens, like Dr *E. coli* bind to bladder and kidney cells under the influence of hormones, thus any acute changes in host hormones like estrogen or progesterone increase the susceptibility of women to UTIs by these bacteria ^1, 16, 17, 44^. Estrogen has immunomodulatory effects on the epithelium and other immune cells mediating actions via ER subtypes to finally affect expression of various innate immune factors. Varying tissue specific distribution of the ER subtypes has been reported ^45-51^, including the bladder ^52-56^. Nuclear ER subtypes bind to estrogen responsive elements (EREs) in the promoter region of cytokines to induce cytokine gene transcription, thereby modulating the inflammation ^37, 51^. However, the contributions of ER subtypes in regulating the bacterial colonization or regulation of pro-inflammatory or anti-inflammatory responses in the bladder for affecting the bacterial colonization or clearance have not been studied.

The objective of the current study was to establish an *in vitro* UTI model in HBEC 5637 cells to understand the involvement of ER subtypes in modulating early innate immune responses and regulating Dr *E. coli* colonization and infection outcome. The presence of Dr *E. coli* colonizing factor CD55 was confirmed on these cells to ensure effective colonization by these bacteria. In addition, the presence of all three ER subtypes was also confirmed on HBEC 5637 cells. Among the ERs, the relative expression of GPR30 was the highest followed by ERβ when compared to ERα. This is the first study reporting the expression of GPR30 and Dr *E. coli* colonization receptor, CD55 on HBEC 5637 cells. We also confirmed that the presence of both host cell CD55 and bacterial Dr fimbriae were required for Dr *E. coli* invasion in HBEC cells.

ER subtype modulation by specific agonist or antagonist treatment of HBEC 5637 cells resulted in differential expression of bacterial colonization receptor CD55, impacting the invasion by Dr *E. coli* in these cells. The hormonal drug effects on these cells also led to differential *tnfa* and *il10* gene expression. Activation of ERα by PPT in 5637 cells resulted in significantly reduced bacterial invasion by 70-85%, and this protection was partially reversed by ERα antagonist MPP. The PPT-mediated protection against bacterial invasion can be explained by the observed downregulation of CD55 and IL10 and the upregulation of TNFα gene expression in PPT treated uninfected cells. In contrast, suppression of ERα in bladder cells by MPP led to upregulation of CD55 and IL10 and downregulation of TNFα gene expression. These results are in contrast to our observations made in the mouse bladder in our *in vivo* UTI model study ^36^, where MPP drug treatment showed relatively more protection against Dr *E. coli* invasion in the bladder than PPT. These protective effects of MPP in the mouse bladder were mediated by downregulation of CD55 and upregulation of TNFα expression. The conflicting results observed in mouse bladder tissue versus human bladder cells may be attributed to variability of distribution and action of ER subtypes in the bladder of two species. Further, the gene expression patterns of TNFα and IL10 were found to be reversed in PPT treated infected cells. The opposing gene expression of these two pro and anti-inflammatory cytokines may be the result of bladder cells undergoing the bacterial clearance and also the subsequent restoration to the homeostasis state. CD55 expression in the infected cells was not determined as CD55 is often cleaved off from the infected cells.

ERβ activation in the bladder cells by DPN significantly reduced Dr *E. coli* invasion by 30-70%, however to a lesser extent than observed with ERα agonist, PPT. DPN-induced protection in these bladder cells was completely reversed by ERβ antagonist, PHTPP. The reduced bacterial invasion observed in DPN treated cells was facilitated by downregulation of CD55 and IL10 and upregulation of TNFα gene expression observed in the uninfected state. However, the suppression of ERβ by PHTPP treatment in these cells resulted in the upregulation of CD55 and IL10 gene expression at some of the PHTPP doses and downregulation of TNFα gene expression in the uninfected state, that led to an increased bacterial invasion as observed in the infected cells.

In contrast to the actions of ERα and ERβ agonists, GPR30 agonist, G1 treatment in HBEC 5637 cells increased CD55 and IL10 gene expression but decreased TNFα expression in uninfected cells, leading to high bacterial invasion by Dr *E. coli*. To our surprise, blocking of GPR30 receptor by its antagonist G15 reduced CD55 and IL10 gene expression but increased the TNFα expression in the bladder cells, which reduced Dr *E. coli* invasion by 20-50% in the infected cells. However, this protection mediated by G15 against bacterial invasion was lesser than the protection mediated by ERα agonist, PPT. The TNFα and IL10 gene expression patterns were reversed in G15 treated infected cells that showed efficient bacterial clearance and restoration of homeostasis.

In summary, our results show that the nuclear estrogen receptors, ERα and ERβ mediate protection against Dr *E. coli* invasion in HBEC cells by downregulating CD55 and IL10 but upregulating TNFα gene expression. Further, we also demonstrate an opposing role played by the membrane receptor, GPR30 in these bladder cells with respect to the other two ER subtypes in mediating protection against Dr *E. coli* invasion by modulating the innate immune responses. Our siRNA transfection studies in HBEC 5637 cells further confirmed the protective role of ERα against bacterial invasion and its immunomodulatory effects on innate immune responses. Silencing of ERα by siRNA in HBEC 5637 cells led to significantly high CD55 and IL10 gene expression and suppressed pro-inflammatory TNFα gene expression which were responsible for the observed high bacterial invasion in *esr1*^*-/-*^ HBEC 5637 cells, further confirming the protective role of ERα in bladder cells. However, ERα antagonist, MPP induced only a partial protection and modulation of TNFα and IL10 gene expression in HBEC 5637 cells compared to PPT and that may be due to the limitation of the doses selected for the experiments.

These diverse immunomodulatory effects of the ER subtypes in the bladder cells may be a result of their differential gene transcription and signaling mechanisms at the cellular level, thus affecting the innate immunity ^37^. Though ERα and ERβ are mostly known to have antagonistic effects on each other ^57-60^, some studies have shown that they cross-talk to produce a cumulative effect as observed in our studies ^61-64^. Depending on the cell type and the inflammatory milieu, these nuclear receptors can have both pro- and anti-inflammatory effects ^40, 65-68^, by modulating the expression of important cytokines like TNFα. The anti-inflammatory effects that were observed in our study as a result of GPR30 activation by G1 have also been reported in two separate *in vivo* studies conducted in male mice and diabetic rats where production of IL10 was observed affecting the T-helper 17 cells ^69, 70^ Further, our results with G15 antagonist were supported in the recent *in vitro* and *in vivo* mouse brain studies. Pro-inflammatory responses were observed in rat microglial cells treated with G15 or by siRNA-mediated *gper1* gene knockdown ^71^ and also in GPR30 knockout mice ^72^. Similarly, in our recent studies in human A431 cuteneous squamous carcinoma cells, the agonist-based activation of ERα, ERβ and GPR30 differentially upregulated CD55 expression that was downregulated by actions of their specific antagonists ^73^.

Some of the recent studies have reported the possibility of cross-talk between GPR30 and the nuclear ERs (ERα and ERβ) in various cell types such as human breast cancer cells and monocytes ^74-76^. Other studies have reported that GPR30 can either have compensatory effects for the absence of nuclear ERs or antagonistic effects in the presence of nuclear ER subtypes as observed in nuclear ER positive and negative breast cancer cell lines ^77, 78^.

## 5. Conclusions

In summary, this is the first study reporting the differential role of nuclear versus membrane ER subtypes in mediating protection against Dr *E. coli* invasion in human bladder cells via modulation of innate immune molecules, CD55, TNFα and IL10 expression. Estradiol regulation of TNFα expression has been reported in ER subtype positive breast cancer cells ^79, 80^. Further, we also report for the first time the ER subtype-mediated inverse regulation of CD55 and TNFα but a direct regulation of CD55 and IL10 gene expression in uninfected bladder cells under the homeostasis state. To maintain the homeostasis state and to avoid hyper activation or suppression of inflammation at the barrier sites in healthy individuals, it is crucial to regulate the expression of TNFα and IL10 ^25^. These opposite trends in TNFα and IL10 expression have also been reported in the gastrointestinal tissues of normal and Irritable Bowel Syndrome patients showing the variable expression levels of ER subtypes ^35^.

Further studies are needed to investigate the ER-regulated downstream signaling in human bladder epithelial cells that result in boosting the innate immune responses and expelling uropathogens.

Our results may have important clinical significance for discovering new ER-subtype based approaches for generating innate immune responses in the bladder. In addition, the ER subtypes can serve as future targets for developing various novel therapies for mediating bacterial clearance in the bladder by boosting the innate immune responses against recurrent UTIs. In the absence of any viable vaccine and the rising number of multi-drug resistant UPEC strains, such novel therapies may have a huge potential for bringing a paradigm shift towards the development of non-antibiotic based therapies that are highly needed for dealing with the pathogenesis of UTIs.

## Acknowledgements

We would also like to thank Batuel Okda for technical assistance in cell culture experiments. The authors would like to specially thank Dr. Subhas Das for sharing his technical expertise in molecular biology.

